# Disentangling Arousal and Attentional Contributions to Pupil Size: Toward Simultaneous Estimation of Emotional and Cognitive States

**DOI:** 10.1101/2025.04.22.649972

**Authors:** Taihei Tsutsumi, Kei Kanari

**Affiliations:** Division of Integrated Science and Agriculture, Graduate School of Regional Development and Creativity, Utsunomiya University, Utsunomiya, Tochigi, Japan; Advanced Institute of So-Go-Chi (Convergence Knowledge) Informatics, Tohoku University, Sendai, Miyagi, Japan; Department of Information and Communications Engineering, Institute of Science Tokyo, Yokohama, Kanagawa, Japan

## Abstract

Pupillary response serves as a noninvasive physiological marker of autonomic activity, modulated by both emotional arousal and the luminance of attended visual stimuli. However, these factors often co-occur, making it difficult to disentangle their individual contributions. This study investigated how auditory emotional stimuli and attention-guided changes in luminance independently influence pupil dynamics. Participants were instructed to direct their visual attention to either white or black moving dot patterns, and to shift their attention upon the onset of an emotional sound. Pupil responses were z-normalized on a per-trial basis, and optokinetic nystagmus (OKN) was recorded to verify attentional shifts. Results showed that emotional arousal robustly increased pupil size across conditions, while valence had no significant main effect. Pupil size was also modulated by the luminance of the attended stimulus, reflecting parasympathetic influences. No significant interaction between arousal and valence was observed, suggesting additive rather than interactive effects. These findings emphasize the methodological necessity of controlling attentional luminance in pupillometric studies and highlight the potential of pupil-based measures for emotion-aware system design. Crucially, because pupil dynamics reflect both luminance-guided attention and emotional arousal via distinct autonomic pathways, combining pupillometry with nonverbal indicators of attention—such as optokinetic nystagmus (OKN)—may enable the simultaneous estimation of attentional focus and emotional state in real time.

## Introduction

In recent years, advances in artificial intelligence (AI) have sparked growing interest in the estimation of psychological states based on physiological signals. Applications extend beyond medicine [1] and autonomous driving [2] to the development of real-time, objective emotion recognition systems through AI-based human sensing technologies [3]. Among the various physiological indices available, the pupillary response has emerged as a particularly promising measure. It offers a noninvasive and automatic window into internal states such as emotion and attention [4].

Pupil diameter is regulated by the interplay between the sympathetic and parasympathetic branches of the autonomic nervous system. Sympathetic activation leads to pupil dilation, while parasympathetic activation causes constriction [5,6]. As a result, the pupillary response is considered a sensitive indicator of emotional arousal and changes in vigilance, and it has been widely utilized in psychology, neuroscience, and human–computer interaction.

However, pupil size is influenced not only by internal psychological states but also by external visual factors such as ambient luminance and visual attention. These confounding effects complicate the interpretation of pupil diameter as a pure index of emotion. Notably, previous research has shown that pupil size automatically adjusts in accordance with the luminance of the attended region of the visual field [7–9], which may interact with emotion-induced pupil dilation. Ignoring such interactions can lead to erroneous conclusions about emotional processing.

To overcome these challenges, recent studies have sought to disentangle sympathetic and parasympathetic contributions by manipulating visual background luminance [10–12]. For instance, Wardhani et al. [11] demonstrated that emotional sounds presented against high-luminance backgrounds influenced both subjective ratings and pupil size, emphasizing the role of environmental context. Cherng et al. [12] further reported that emotionally evoked pupil dilation and saccade-related pupil changes originate from distinct autonomic pathways—sympathetic and parasympathetic, respectively.

Yet, these studies typically maintained constant background luminance throughout each trial, even before and after the emotional stimulus onset. Therefore, it remains unclear how pupil responses are modulated under more naturalistic conditions, where an emotional sound not only captures attention but also coincides with dynamic changes in the luminance of the attended visual field.

To address this gap, the present study concurrently presented visual stimuli with differing luminance and emotional sounds, requiring participants to shift visual attention from one luminance level to another. We further evaluated the efficacy of the attentional manipulation by analyzing optokinetic nystagmus (OKN), a reflexive eye movement induced by motion stimuli that has recently been recognized as a robust marker of selective visual attention [13–15].

Accordingly, the primary goal of this study was to systematically disentangle the independent and interactive effects of auditory-induced emotional arousal and luminance-guided visual attention on pupil responses. By concurrently manipulating emotional sounds and luminance-directed attention—and by employing OKN as an objective marker of attention shifts—we aimed to clarify the distinct autonomic pathways underlying pupil dynamics and to explore their potential applications in real-time, emotion-aware systems.

## Methods

### Participants

The study was conducted in accordance with the Declaration of Helsinki and was approved by the Ethical Review Committee on Research Involving Human Subjects at Utsunomiya University (protocol code: H21-0010; approved on April 21, 2021). Participants were recruited between May 25, 2021, and October 21, 2021, following ethical approval from the institutional review board. A total of 22 Japanese university students (aged 18–24 years; 14 females; M = 20.9, SD = 1.7) took part in the experiment. All participants were adults (18 years or older), and no minors were included in the study. All had normal or corrected-to-normal vision, with a visual acuity of 1.0 or higher. Written informed consent was obtained from all participants prior to their participation.

### Apparatus

The experiment was conducted in a dark room. Visual stimuli were displayed on a liquid crystal display monitor (ColorEdge CS2420-ZBK, EIZO Corporation, Ishikawa, Japan) with a resolution of 1920 × 1200 pixels, a refresh rate of 60 Hz, and a screen size subtending a visual angle of 48.9° × 31.7°. Eye movements and pupil size were recorded at 500 Hz using a video-based eye tracker (iRecHS2 Ver. 660) [16].

Stimulus presentation and timing control were implemented using MATLAB R2017b (MathWorks, Natick, MA, USA) with the Psychophysics Toolbox [17], running on a MacBook Pro (Apple Inc., Cupertino, CA, USA) under macOS Sierra 10.12.6. The viewing distance was fixed at 57 cm using a chinrest. Auditory stimuli were presented monaurally via wired earphones.

### Stimuli

An example of the visual stimuli is shown in Fig. 1. The stimuli consisted of moving random dot patterns that differed in luminance (white or black) and always moved in opposite directions (leftward or rightward). The entire pattern was confined to a circular area with a radius of 10.57° of visual angle and moved at a constant speed of 16.4 deg/s. Each dot had a diameter of 0.27°. The luminance levels were 222.33 cd/m^2^ for white dots, 0.26 cd/m^2^ for black dots, and 49.27 cd/m^2^ for the gray background.

**Fig 1.**
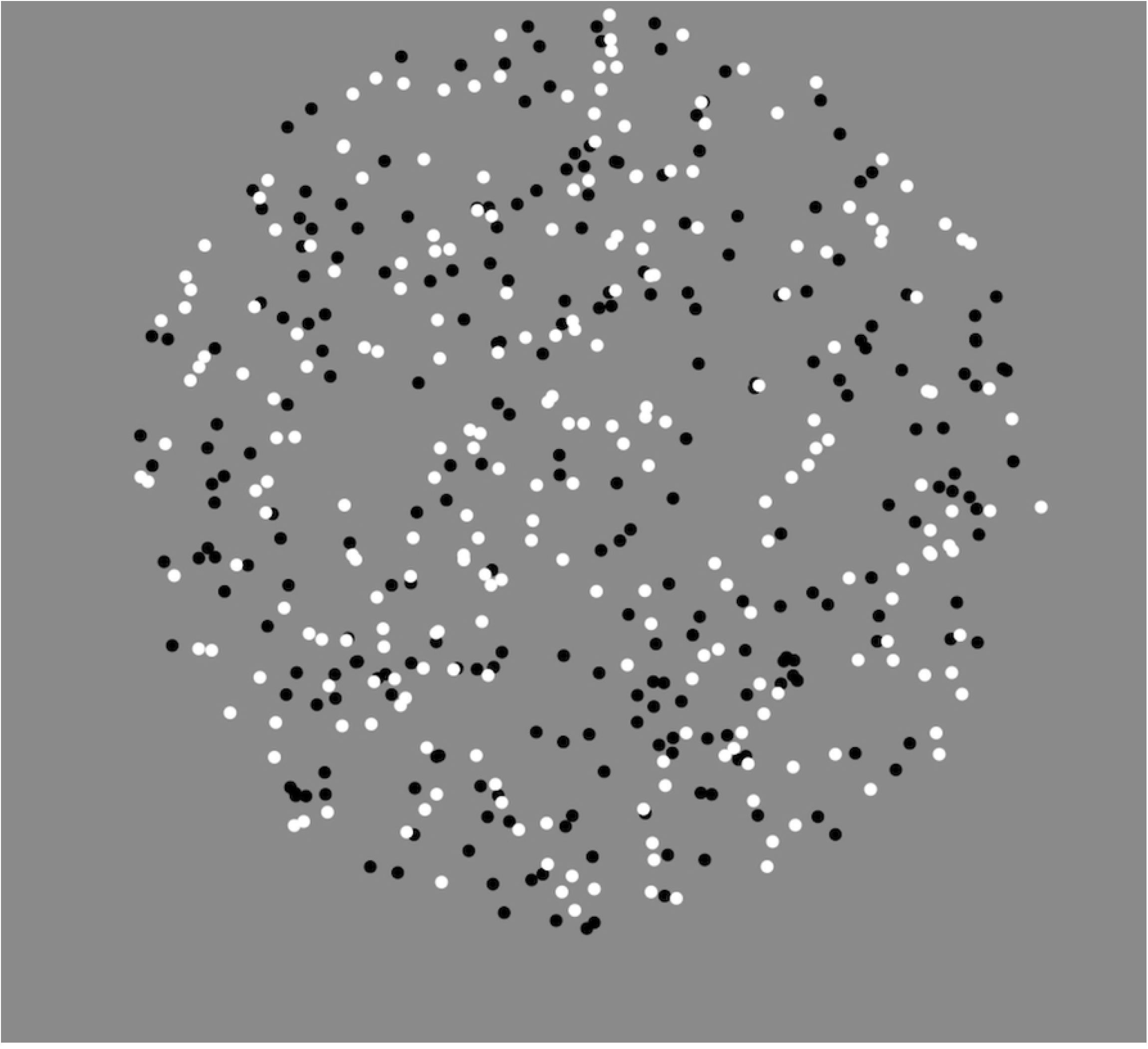
Example of the visual stimuli used in the experiment. The stimulus consisted of a random dot pattern composed of white and black dots presented on a gray background. In each condition, the white and black dots moved in opposite directions (leftward or rightward) as a whole pattern. The entire pattern was constrained to move within a circular area with a radius of 10.57° of visual angle.

Auditory stimuli were selected from the Expanded Version of the International Affective Digitized Sounds (IADS-E) [18], developed by Yang et al. (2018) as an extension of the IADS-2 system [19]. The IADS-E comprises 935 digitally recorded, naturalistic sound clips that reflect common experiences in daily life, such as babies crying, footsteps, thunder, and background music [20]. Each sound is characterized by three Self-Assessment Manikin ratings—Valence, Arousal, and Dominance [21]— as well as by basic emotional categories (e.g., happiness, sadness, fear). We selected 20 arousal-varied sounds and 20 valence-varied sounds (40 in total), ensuring that the rating distributions within each set were uniformly spread across the 1–9 scale. The Arousal and Valence sets were mutually exclusive.

### Procedure

A schematic of the trial sequence is shown in Fig. 2. At the beginning of each trial, participants received instructions regarding which dot luminance (white or black) to attend to, both before and after the auditory stimulus. After pressing a key, a fixation point appeared at the center of the screen against a gray background for 4–5 seconds. The fixation point was presented in the color of the dot to which participants were initially instructed to attend, and a text label indicating the initial target dot was also displayed above it. The duration of the fixation period was jittered to prevent participants from anticipating the onset of the visual stimulus.

**Fig 2.**
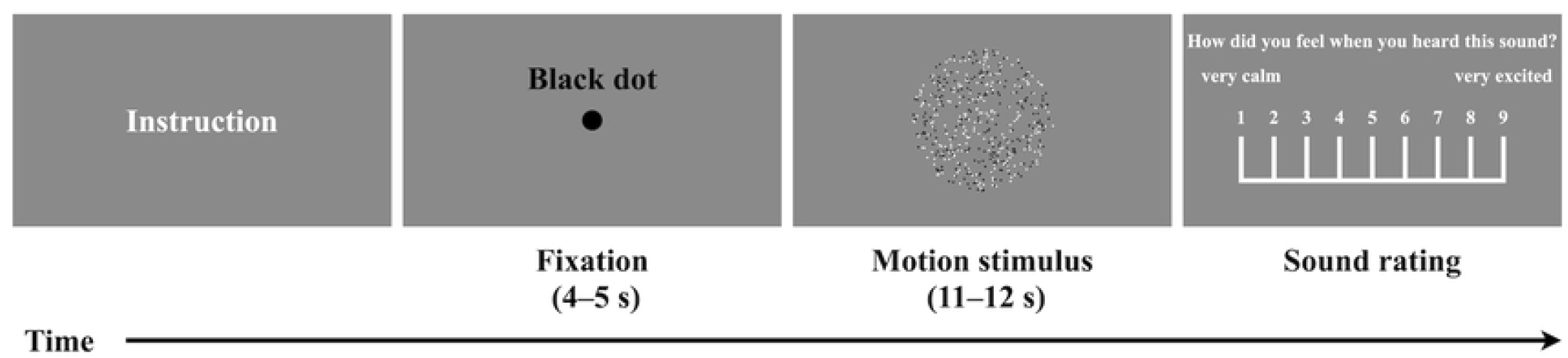
Schematic of the trial sequence. Each trial began with an instruction screen indicating the target dot luminance (black or white) to which participants were instructed to attend. This was followed by a fixation period (4.0–5.0 s), a stimulus presentation phase (11.0–12.0 s) during which moving random dots were displayed, and finally a sound rating phase, during which participants evaluated the auditory stimulus in terms of arousal and valence.

Subsequently, the random dot patterns were presented for 11–12 seconds, with white and black dots moving in opposite directions. An emotional sound was played 5–6 seconds after motion onset, with the timing jittered to avoid predictability. The sound lasted for 6 seconds. Participants were instructed to shift their attention to the dot pattern of the opposite luminance immediately upon hearing the sound.

Following each trial, participants evaluated the auditory stimulus on two dimensions using 9-point scales: arousal was rated first, followed by valence. For Arousal, 1 indicated “very calm” and 9 “very excited”; for Valence, 1 indicated “very negative” and 9 “very positive.”

Two experimental conditions were used: the BW condition, in which attention shifted from black to white dots, and the WB condition, in which attention shifted from white to black. Each condition consisted of 40 trials (one per sound), resulting in 80 trials in total. The order of sound presentation was randomized for each participant. Breaks were provided between conditions and could also be taken between trials as needed.

### Eye movement and pupil data analysis

In the eye movement analysis, gaze position values exceeding 100,000 were treated as missing. The analysis window included 4 seconds before and 6 seconds during the sound presentation (total: 10 seconds). Outliers, defined as values more than three standard deviations from the mean, were replaced with the nearest valid data points. OKN slow phases were then extracted. Horizontal eye velocity (in deg/s) was computed via differentiation of gaze position, and segments exceeding 16.4 deg/s were excluded to remove saccades and fast OKN components. The data were then smoothed using a Gaussian-weighted moving average with a 1000-ms window.

For pupil data analysis, blinks were detected and removed using the default algorithm of the iRecHS2 system. Outlier processing followed the same procedure as for gaze data. Pupil size was smoothed using a 400-ms Gaussian filter. Within each trial, pupil values from –5 to +6 seconds relative to sound onset were z-scored. The average pupil size during the 6-second auditory period was then used as the dependent variable in subsequent regression analyses.

## Results

### Time course of pupil and OKN changes

Figure 3 illustrates the time course of average pupil size across participants, with time 0 indicating the onset of the auditory stimulus. Pupil dilation was observed after sound onset in both the BW and WB conditions. Subsequently, a constriction response occurred in the BW condition in response to the white stimulus, while further dilation was observed in the WB condition in response to the black stimulus. Statistical analysis followed the method described by Einhäuser et al. [22], using point-wise *t*-tests and controlling the false discovery rate (FDR) with the Benjamini–Hochberg procedure [23]. Time points exceeding the FDR threshold (*p*_thresh_ = 0.047) are indicated by black horizontal lines in Fig. 3.

**Fig 3.**
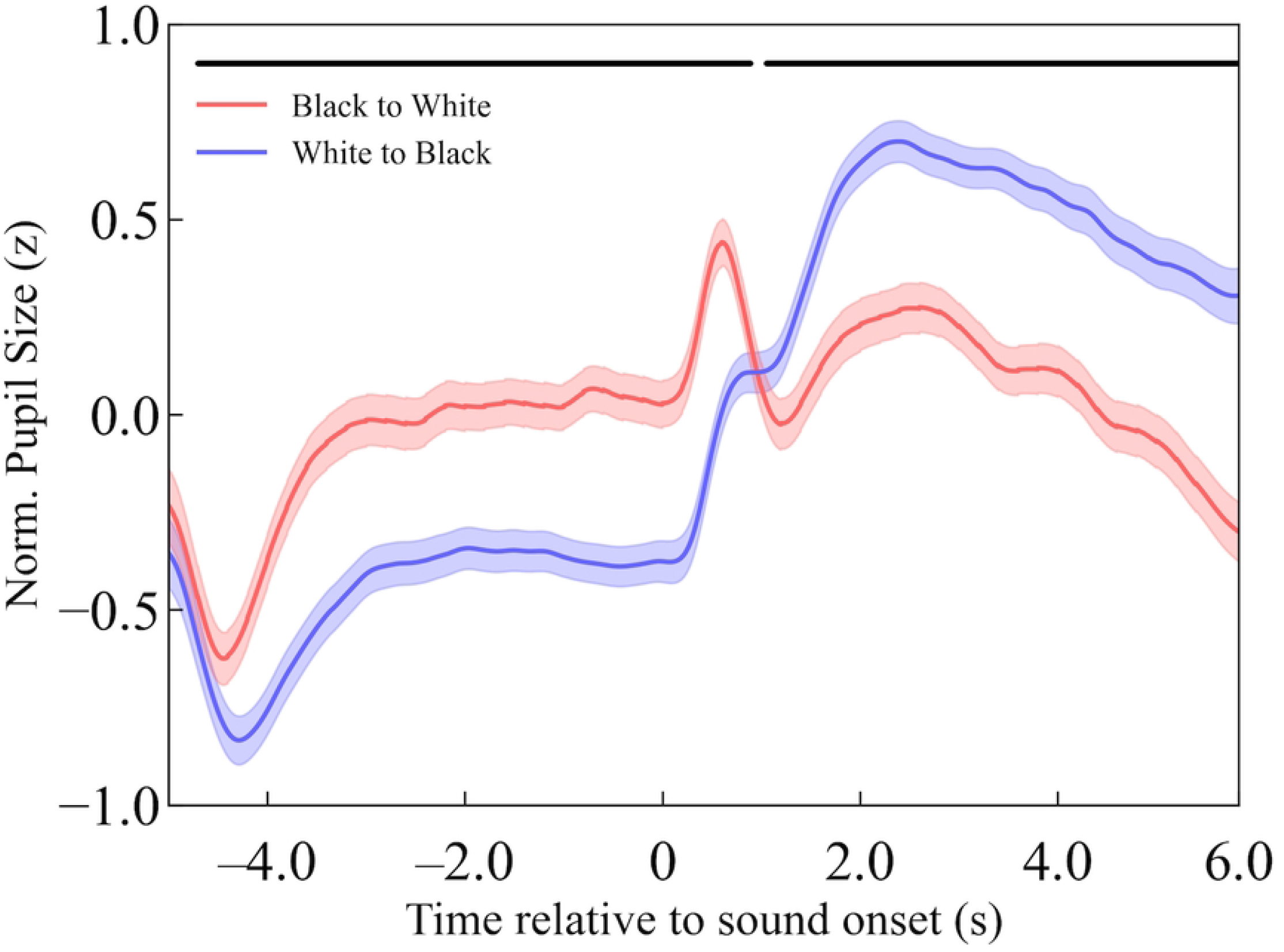
Time course of pupillary responses before and after auditory stimulus onset. The x-axis represents time (in seconds), with 0 indicating the onset of the auditory stimulus. The y-axis shows pupil size in z-scored units, calculated from −5 to +6 seconds relative to sound onset within each trial. The red line indicates the BW condition (attention shifted from black to white), and the blue line indicates the WB condition (white to black). Shaded areas represent 95% confidence intervals. Black horizontal lines indicate time points at which statistical significance was reached following FDR correction (*p*_thresh_ = 0.047).

Figure 4 shows the time course of OKN slow-phase velocity across all participants, used to evaluate the effectiveness of the attentional manipulation. Before sound onset, OKN slow-phase velocity aligned with the motion direction of the initially attended stimulus. Approximately 0.5 seconds after sound onset, the velocity shifted in the opposite direction, consistent with attention switching. Black horizontal lines in Fig. 4 indicate time intervals during which statistically significant differences were observed (*t*-test, FDR-corrected; *p*_thresh_ = 0.047).

**Fig 4.**
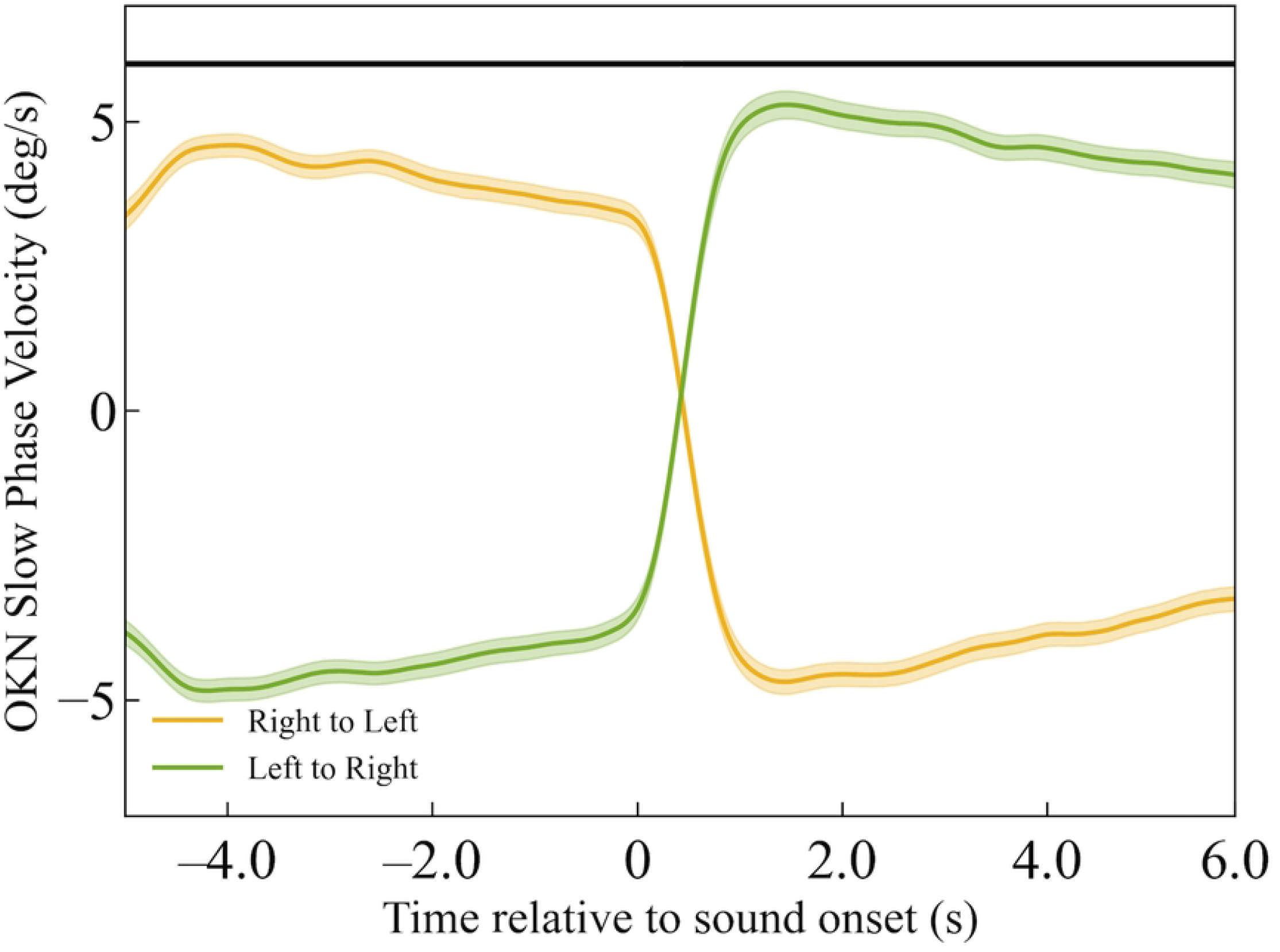
Time course of OKN slow-phase velocity before and after auditory stimulus onset. The vertical axis indicates the OKN slow-phase velocity (deg/s), with positive values representing rightward eye movements relative to the display. The green line shows the average data for the condition in which attention shifted from leftward-moving dots to rightward-moving dots, while the yellow line shows the condition with the opposite shift (right to left). Shaded areas represent 95% confidence intervals. Black horizontal lines indicate time points at which statistical significance was reached following FDR correction (*p*_thresh_ = 0.047).

### Regression analysis: predicting pupil response from emotional ratings

To examine how well Arousal and Valence predicted pupil responses, a multiple regression analysis was conducted. In the BW condition, the coefficient of determination (*R*^2^) was 0.66. Arousal showed a significant positive regression coefficient (unstandardized *β* = 0.083, 95% CI [0.058, 0.108], *p* < .001; standardized *β* = 0.846, 95% CI [0.589, 1.102], *p* < .001), whereas Valence had no significant effect (unstandardized *β* = 0.006, 95% CI [–0.024, 0.037], *p* = .68; standardized *β* = 0.05, 95% CI [–0.204, 0.309], *p* = .68). In the WB condition, *R*^2^ was 0.46. Arousal again showed a significant positive effect (unstandardized *β* = 0.045, 95% CI [0.025, 0.064], *p* < .001; standardized *β* = 0.76, 95% CI [0.431, 1.098], *p* < .001), while Valence remained non-significant (unstandardized *β* = 0.009, 95% CI [–0.013, 0.031], *p* = .41; standardized *β* = 0.138, 95% CI [–0.194, 0.471], *p* = .41).

In the BW condition, Arousal ratings were significantly positively correlated with pupil responses, as indicated by Pearson’s correlation coefficient (*r* = 0.81, 95% CI [0.67, 0.90], *p* < .001). In contrast, Valence ratings also showed a significant moderate negative Pearson correlation with pupil responses (*r* = –0.50, 95% CI [–0.70, –0.22], *p* = .001). In the WB condition, Arousal ratings were significantly positively correlated with pupil responses (*r* = 0.67, 95% CI [0.45, 0.81], *p* < .001), whereas Valence ratings showed a significant but weaker negative Pearson correlation (*r* = –0.38, 95% CI [–0.62, –0.08], *p* = .016). Figures 5 and 6 present the relationships between pupil responses and emotional ratings under the BW (black-to-white) and WB (white-to-black) conditions, respectively.

**Fig 5.**
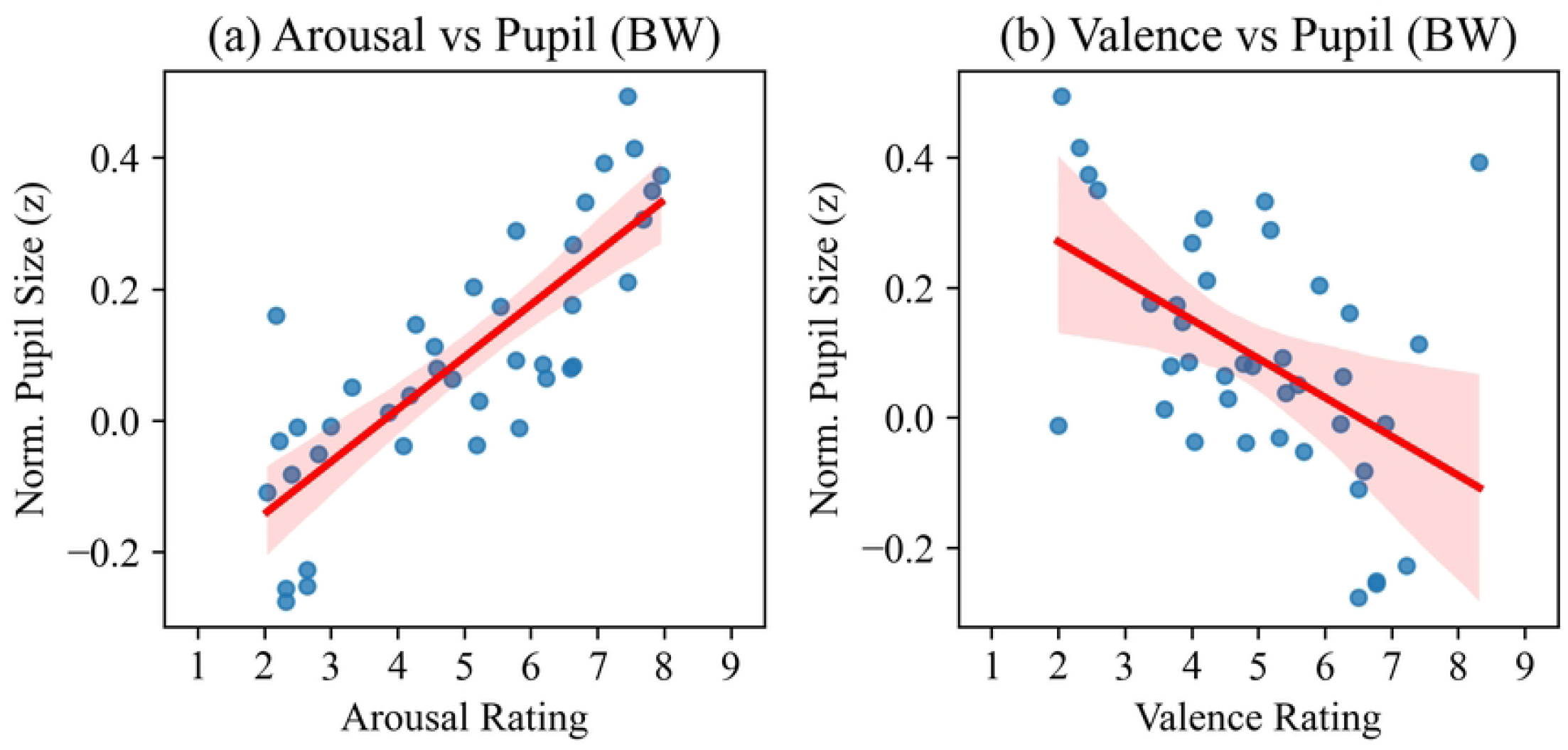
Relationships between pupil responses and emotional ratings under the BW (black-to-white) condition. (a) Relationship between pupil responses and arousal ratings. Arousal ratings showed a statistically significant positive Pearson correlation with pupil responses (*r* = .81, 95% CI [0.67, 0.90], *p* < .001). (b) Relationship between pupil responses and valence ratings. Valence ratings showed a statistically significant negative Pearson correlation with pupil responses (*r* = –.50, 95% CI [–0.70, –0.22], *p* = .001). In each scatter plot, each dot corresponds to one of the 40 auditory stimuli. The red line represents the linear regression line fitted to the data, and the surrounding shaded area indicates the 95% confidence interval of the regression estimate.

**Fig 6.**
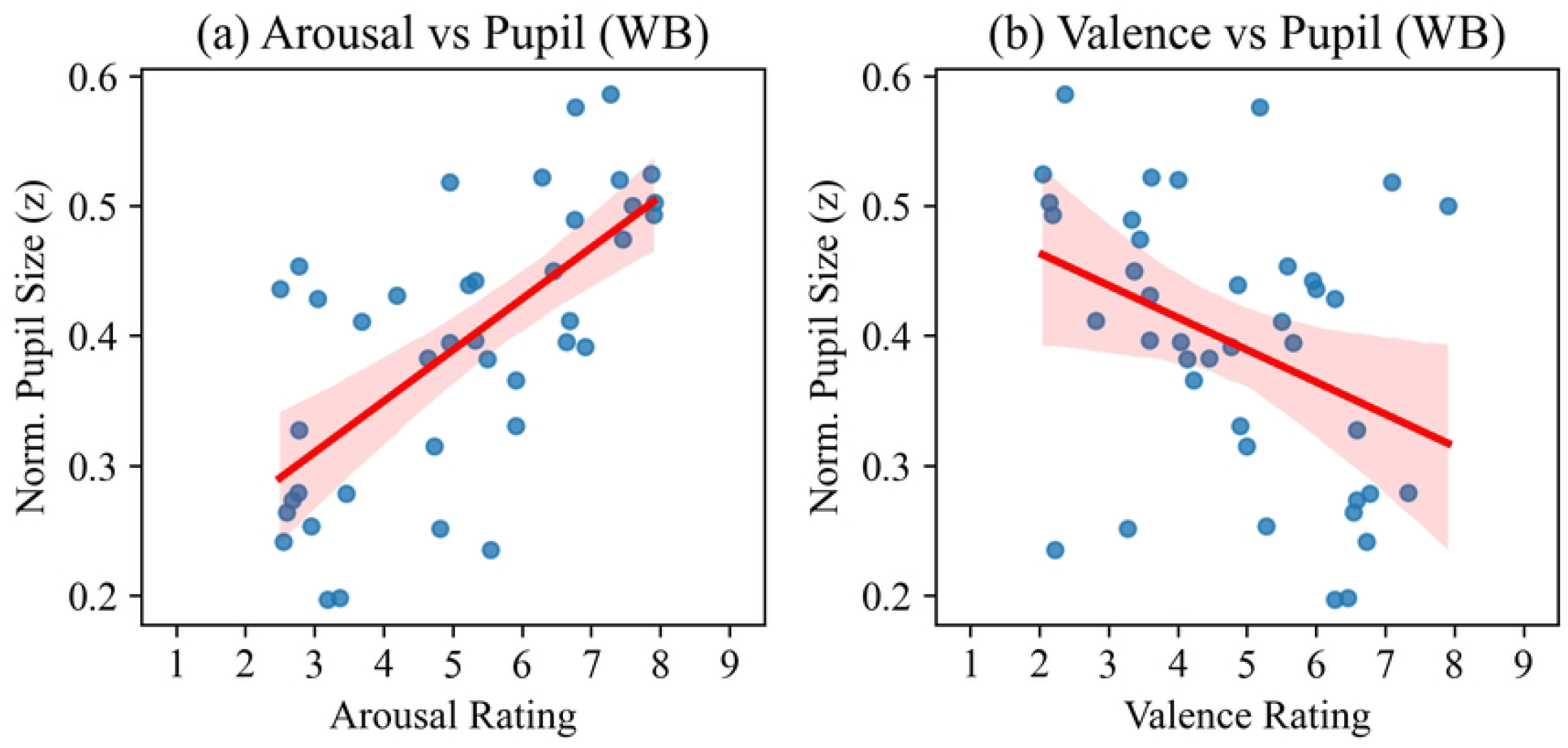
Relationships between pupil responses and emotional ratings under the WB (white-to-black) condition. (a) Relationship between pupil responses and arousal ratings. Arousal ratings showed a statistically significant positive Pearson correlation with pupil responses (*r* = .67, 95% CI [0.45, 0.81], *p* < .001). (b) Relationship between pupil responses and valence ratings. Valence ratings also showed a statistically significant negative Pearson correlation with pupil responses (*r* = –.38, 95% CI [–0.62, –0.08], *p* = .016).

### Multicollinearity assessment (VIF)

To evaluate potential multicollinearity between Arousal and Valence, variance inflation factors (VIFs) were calculated. VIF values were approximately 3.1 for both Arousal and Valence (BW: 3.07; WB: 3.13), indicating no substantial multicollinearity. The Pearson correlation coefficients between arousal and valence were moderately negative and statistically significant in both conditions (BW: *r* = –0.65, 95% CI [–0.80, –0.43], *p* < .001; WB: *r* = –0.68, 95% CI [–0.82, –0.47], *p* < .001).

### Regression Diagnostics

To assess the assumptions of the regression models, residual plots, Q–Q plots, and Shapiro–Wilk tests were conducted. Residual and Q–Q plots for both conditions are presented in S1 Fig and S2 Fig. Residuals were randomly distributed with respect to the fitted values, and the LOWESS trendlines were generally flat, supporting the assumptions of linearity and homoscedasticity. The Q–Q plots indicated that residuals closely followed the expected normal distribution, and the Shapiro– Wilk tests did not reject the null hypothesis of normality (BW: *p* = .054; WB: *p* = .655). These results suggest that the basic assumptions of linear regression were reasonably satisfied in both conditions.

### Interaction effects of arousal and valence

Including the Arousal × Valence interaction term in the mixed-effects regression models did not reveal a significant effect in either condition. In the BW condition, the interaction was not statistically significant (unstandardized *β* = –0.001, 95% CI [–0.015, 0.012], *p* = .830), nor was it in the WB condition (unstandardized *β* = –0.002, 95% CI [–0.012, 0.008], *p* = .673).

### Comparison between conditions (BW vs. WB)

The BW (black-to-white) condition demonstrated greater explanatory power (*R*^2^ = 0.66) than the WB (white-to-black) condition (*R*^2^ = 0.46), particularly with respect to the effect of arousal, which was statistically significant only in the BW condition. In contrast, Valence and the Arousal × Valence interaction did not show significant effects in either condition. These results indicate that arousal had a stronger and more consistent effect on pupil responses, particularly in the BW condition.

## Discussion

### Arousal dominates over valence in predicting autonomic pupil response

This study demonstrated that subjective arousal ratings exerted a consistent and significant positive effect on pupil dilation in both the BW and WB conditions. This finding aligns with prior research suggesting that increased arousal enhances sympathetic activity, thereby resulting in pupil enlargement [11,12,24]. Notably, the high coefficient of determination observed in the BW condition (*R*^2^ = 0.66) indicates that emotional arousal was robustly reflected in pupillary dynamics.

In contrast, valence ratings did not yield significant effects in the multiple regression models for either condition, suggesting a limited independent contribution to pupil responses. However, Pearson correlation analyses revealed significant negative associations between valence and pupil size, particularly in the BW condition (*r* = –0.50, *p* = .001). This apparent discrepancy may be explained by the moderate negative correlation observed between arousal and valence (BW: *r* = –0.65; WB: *r* = –0.68). As a result, valence may share predictive variance with arousal, such that its effect is statistically accounted for when arousal is included in the regression model. In this view, valence could influence pupil responses indirectly, or through shared variance with arousal, rather than exerting an autonomous effect.

These findings are consistent with previous studies reporting a stronger association between pupil size and arousal than between pupil size and valence [24,25]. They further support the notion that pupillary responses are primarily modulated by the intensity of emotional arousal, which more directly engages sympathetic pathways, whereas valence—reflecting emotional pleasantness—may play a subordinate or context-dependent role in autonomic modulation.

These results are also compatible with dimensional models of emotion, which posit that arousal and valence represent orthogonal dimensions of affect and may contribute independently to autonomic responses [26,27]. In this framework, arousal is typically associated with sympathetic nervous system activation, leading to increased pupil dilation, while valence may exert a more limited or indirect influence. The present findings support this distinction: arousal robustly predicted pupil responses across conditions, whereas valence, despite showing some correlation with pupil size, did not explain additional variance in the regression models. This suggests that autonomic modulation of pupil diameter primarily reflects the intensity of emotional activation, rather than its hedonic quality.

### Additive rather than interactive effects of emotional dimensions

To further examine whether arousal and valence interact in modulating pupil responses, we included an Arousal × Valence interaction term in the regression model. The interaction effect was not statistically significant in either the BW or WB condition, suggesting that arousal and valence operate in an additive rather than interactive manner. In other words, highly arousing stimuli consistently enhanced pupil dilation regardless of valence, and valence alone did not modulate the effect of arousal. These findings are compatible with dimensional models of emotion in which arousal and valence contribute separately to affective processing and autonomic responses [26,27].

### Condition differences (BW vs. WB): attenuation and asymmetry

The observed difference in explanatory power between the BW and WB conditions suggests that the temporal dynamics of neural responses associated with attentional shifts may vary across conditions. Specifically, in the WB condition, attention was initially directed toward a bright stimulus (white dots), likely enhancing parasympathetic activity and inducing pupil constriction. This constriction temporally overlapped with sympathetic activation elicited by emotional arousal from the auditory stimulus, resulting in an attenuated overall pupil response and a relatively weaker effect of arousal. Such opposing autonomic influences are consistent with the integrative regulatory functions of the locus coeruleus (LC), which coordinates both sympathetic and parasympathetic inputs to modulate pupil diameter [25].

### Temporal dynamics of pupil response: emotional and attentional interplay

Time-resolved analysis of pupil responses revealed a brief constriction between –5 and –4 seconds, likely reflecting reflexive parasympathetic activation at the onset of the visual stimulus. Between –4 and 0 seconds, sustained parasympathetic activity appeared to correspond to the luminance of the initially attended stimulus. Approximately 0.5 seconds after auditory onset, a second phase of parasympathetic modulation emerged, now aligned with the newly attended stimulus. These temporal patterns suggest that sympathetic activation driven by emotional arousal and parasympathetic modulation driven by attentional luminance cues alternated over time, underscoring a dynamic interplay between emotional and attentional processes.

### Objective validation of attention manipulation: evidence from OKN

The effectiveness of the attentional manipulation was objectively validated using OKN slow-phase velocity. Following the auditory cue, the direction of OKN consistently reversed, indicating that participants shifted their attention as instructed. Thus, OKN served as a reliable and nonverbal indicator of attentional control, representing a methodological strength of the present study.

### Statistical validity and autonomic modeling

Variance inflation factors (VIFs) of approximately 3 confirmed that the effects of arousal and valence were assessed independently, with no evidence of problematic multicollinearity, thereby supporting the statistical robustness of the regression models. These results reinforce the notion that pupil dynamics emerge from a complex interplay among emotional, attentional, and luminance-driven processes. This interpretation aligns with prior models in which light-induced constriction is primarily mediated by parasympathetic activity via the retino-pretectal pathway, whereas emotional arousal engages sympathetic pathways involving the amygdala, hypothalamus, and midbrain structures [10,24].

Future research should aim to further elucidate the neural mechanisms underlying these processes—particularly the interactions among the amygdala, thalamus, superior colliculus, and brainstem pupillary control centers—in order to develop more comprehensive models of emotion–attention integration. The finding that valence did not significantly interact with arousal, despite showing a correlation with pupil size, also highlights the need for refined models capable of distinguishing between shared variance and independent effects. These findings offer promising implications for real-time adaptive systems in domains such as driver monitoring and affective computing, where integrated models of emotion and attention could enhance the sensitivity and robustness of human–machine interaction.

### Limitations

This study has several limitations. First, because emotional stimuli and attention-switching instructions were presented simultaneously, their individual contributions to pupil responses could not be temporally disentangled. Given that pupil size reflects both sympathetic dilation and parasympathetic constriction, the effect of arousal may have been underestimated. Future studies should consider temporally separating emotional and attentional cues (e.g., presenting the auditory stimulus prior to the attentional switch).

Second, the primary dependent measure was the mean pupil change during the 6-second auditory period, selected to capture the sustained effects of arousal [28,29]. However, this averaging approach may obscure dynamic characteristics such as onset latency, rise slope, and recovery time. Future research should incorporate time-resolved metrics to more accurately infer underlying psychological states. Analytical approaches such as wavelet decomposition [30] and recurrent neural networks (RNNs) [31] may facilitate the extraction of subtle, nonlinear dynamics from pupil waveforms.

## Conclusion

This study provides empirical evidence that pupillary responses reflect the additive effects of emotional arousal and luminance-guided visual attention, mediated by distinct autonomic pathways. While arousal consistently enhanced pupil dilation, the luminance of the attended stimulus modulated pupil size in a parasympathetically driven manner, independent of emotional valence. The lack of interaction between arousal and valence supports the idea that emotional and attentional processes exert separable influences on the autonomic control of the pupil.

These findings highlight the methodological necessity of accounting for attentional luminance in pupillometric research and suggest that pupil-based measures can serve as robust indicators of emotional arousal under dynamic, attention-involving conditions. Because pupil size reflects both arousal-related sympathetic activation and attention-driven luminance modulation, pupillometry holds promise as a means of simultaneously estimating emotional state and attentional focus from a single physiological signal.

## Acknowledgments

The authors would like to thank all participants for their involvement and cooperation in this study.

## Supporting information

**S1 Fig. Residual plots with LOWESS smoothing lines**. (a) BW condition. (b) WB condition. Each plot shows the residuals from an ordinary least squares (OLS) regression model predicting pupil response from arousal and valence. The horizontal axis represents fitted values; the vertical axis represents raw residuals. A locally weighted scatterplot smoothing (LOWESS) line is overlaid in red. The residuals appear to be approximately randomly scattered around zero without strong patterns, supporting the assumptions of linearity and homoscedasticity.

**S2 Fig. Q–Q plots of standardized residuals for each condition**. (a) BW condition. (b) WB condition. Each plot displays the residuals from an ordinary least squares (OLS) regression model predicting pupil response from arousal and valence. The residuals were standardized and plotted against the theoretical quantiles of the standard normal distribution. The points closely follow the diagonal line, indicating that the residuals are approximately normally distributed in both conditions.

